# Dynamic facial trustworthiness perception in real-time social contexts

**DOI:** 10.1101/2024.08.19.608645

**Authors:** Haoming Qi, Dongcheng He

**Affiliations:** School of Information Network Security, People’s Public Security University of China, Beijing, China; Institute of Dataspace, Hefei Comprehensive National Science Center, Hefei, Anhui, China; Herbert Wertheim School of Optometry and Vision Science, University of California, Berkeley, Berkeley, CA, United States

**Keywords:** trustworthiness, EEG, hemispheric asymmetry, affective cognition

## Abstract

Current understanding of the neural mechanisms underlying facial trustworthiness perception is primarily based on studies using static facial stimuli. However, real-life social interactions are dynamic and complex, and the neural processes involved in such naturalistic contexts remain largely unexplored. In this study, we analyzed EEG data collected during a deception game involving two participants: a player and an observer engaged in real-time interaction. The player either followed instructions or made spontaneous decisions to lie or tell the truth, while the observer judged whether to trust the player based solely on their facial expressions. We examined observers’ behavioral data, event-related potentials, and interhemispheric EEG asymmetries in both signal magnitude and instantaneous phase. The results revealed a significant effect of trustworthiness on hemispheric asymmetry in the observer’s centroparietal phase activities between 1500-3000 ms post-stimulus. Subsequent frequency-based analysis revealed that this asymmetry in phase progression was primarily driven by lateralized signal frequency. These findings suggest that the perception of facial trustworthiness involves dynamic hemispheric lateralization. Whereas previous studies using static face stimuli indicate rapid trustworthiness perception, our findings suggest that trustworthiness perception can be modulated by persistent and dynamical affective processing in real-time social contexts.

## 1. Introduction

A line of studies has demonstrated that people rapidly and intuitively form impressions of others’ social attributes, such as trustworthiness, based facial expression perception. In a notable study, Willis & Todorov (2006) examined participants’ judgments of various traits (e.g., trustworthiness, attractiveness, etc.) from face stimuli presented for 100ms, 500ms, and 1000ms. They found that participants’ judgement was highly correlated across all time conditions, suggesting that reliable first impressions regarding the social attributes can be formed in as little as 100ms. Extending this work, Todorov et al. (2009) conducted another experiment with a similar paradigm but employing finer-grained exposure durations ranging from 17 ms to unlimited viewing time. They reported that trustworthiness judgments were significantly above chance after only 33 ms of exposure. Subsequently, using a composite-face paradigm, that aligned trustworthy upper halves of faces and untrustworthy lower halves versus the opposite, Todorov et al. (2010) found that participants failed to discern between these two types of stimuli in judging the trustworthiness with exposure time shorter than 100ms. However, with longer viewing durations, participants rated the composites with trustworthy upper halves more positively, supporting the role of early holistic processing in trustworthiness perception. These studies demonstrated that impressions of social attributes formed from faces are instinctive evaluations, shaped more by perception than by deliberate reasoning. Such findings have been replicated by many subsequent studies (Bar et al., 2006; Porter et al., 2008; Holtz, 2015). Further investigations have explored additional factors influencing the phenomenon of facial trustworthiness perception. For example, Krumhuber et al. (2007) found both genuine and fake smile faces were perceived to be more trustworthy than neutral faces, highlighting the role of emotional cues. Also, Robinson et al. (2014) identified the eyes and mouth as critical facial regions for facial trustworthiness perception, and demonstrated that manipulating the relative saliency of these regions could bias judgments toward trustworthy or untrustworthy. Other factors include the observer’s age (Cassidy et al., 2019), knowledge about others’ social category (Xie et al., 2019; Schmid et al., 2022), experience (Cheung et al., 2024), and even momentary associations (Fenske et al., 2005). These findings emphasize the roles of both perceptual and non-perceptual factors in biasing trustworthiness perception.

EEG has been instrumental in uncovering the temporal dynamics of cortical activities during facial trustworthiness perception, as well as its relationship to other cognitive functions. In an ERP study, Yang et al. (2011) observed a larger C1 component (within 40-90ms post-stimulus) in response to trustworthy compared to untrustworthy faces. This finding supports previous behavioral findings that facial trustworthiness perception occurs at remarkably early stages. Additionally, a larger late positive component was detected in response to trustworthy faces, suggesting increased attention demand. In another study, Calvo et al. (2018) investigated the interaction between emotional expression and trustworthiness perception. They presented faces with multiple types of emotional expression and found that facial expression processing could occur earlier than trustworthiness judgement, potentially reflecting a temporal hierarchy where emotional cues may precede and influence the evaluation of social attributes like trustworthiness. While these studies relied on explicit judgment tasks, other research has employed fast periodic visual stimulation (FPVS) to investigate implicit neural processing of facial trustworthiness. FPVS involves rapidly presenting visual stimuli at fixed frequencies, allowing the extraction of frequency-tagged EEG responses (Rossion, 2014). By updating face stimuli rapidly, this approach allows decoding EEG data patterns induced by relatively automatic and implicit neural functions constrained by a short exposure duration. Using this approach, Swe et al. (2020) presented face stimuli at a base rate of 6 Hz, with trustworthiness systematically varying at 1 Hz. They successfully detected trustworthiness-related neural responses at 1 Hz, providing strong evidence for implicit encoding of facial trustworthiness perception. In another experiment using FPVS, trustworthy faces (oddballs) were interspersed every fifth stimulus among untrustworthy faces, and verse versa, leaving the oddball rate and base rate at 1.2Hz and 6Hz, respectively (Verosky et al., 2020). They found that the trustworthiness of oddball faces had a significant effect on the EEG signals at 1.2Hz. These EEG studies align with behavioral data on the intuitive processing of facial trustworthiness, highlighting the role of implicit mechanisms in trustworthiness perception.

Regarding the cortical regions involved in facial trustworthiness perception, previous studies using fMRI have consistently identified the amygdala, a key region for affective processing, as showing different BOLD signals in response to face stimuli varying in perceived trustworthiness (Engell et al., 2007; Todorov & Engell, 2008; Said et al., 2009; Winston et al., 2013). It is well established that amygdala activity exhibits hemispheric asymmetry, particularly in the context of emotion processing (a review of neuroimaging studies regarding this issue: Baas et al., 2004). Early research using visual masking paradigms indicated that this lateralization depends on the observer’s awareness of emotional facial expressions, with unconscious processing primarily engaging the right amygdala (Morris et al., 1998). Additional theories about the functional differences between the left and right amygdala were also hypothesized, positing that the right amygdala supports rapid, automatic emotional responses, whereas the left amygdala is involved in more sustained and finer modulation of emotional arousal (Gläscher & Adolphs, 2003). Although the exact relationship between amygdala lateralization and EEG measures remains unclear, emotional processing is known to elicit hemispheric asymmetries in EEG signals. For example, the left frontal ERP was found to be lateralized to happy faces, while the right was lateralized to neutral faces (Graham & Cabeza, 2001). Such lateralization effects in EEG signals from emotional valence were also detected based on spectral analysis (Ahern & Schwartz, 1985; Pane et al., 2019) and instantaneous phase analysis (Costa et al., 2006; Val-Calvo et al., 2019; Cao et al., 2020).

Previous research has significantly contributed to our knowledge about the cortical dynamics underlying facial trustworthiness perception and its associated mechanisms. However, the vast majority of these studies have relied on static facial photographs or algorithmically generated face images drawn from databases with preassigned trustworthiness scores. In contrast, real-world social interactions involve dynamic, temporally evolving facial expressions that unfold during interpersonal exchanges. Such naturalistic conditions may modulate trustworthiness perception over time in ways not captured by static stimuli. Yet perceiving actual faces in a real-time manner could complicate the cortical processing, little is known about the neural activities under such conditions. Particularly, the neural signatures revealing trustworthy verses untrustworthy in natural settings may not be readily inferred from previous findings based on static facial stimuli. To address this gap, we analyzed a novel EEG dataset collected simultaneously from a “player” and an “observer” during a task designed to encourage deception. At each time, the player decided to relay either the true or a false information to the observer, who viewed the player’s face before choosing to trust the player or not. This design enabled us to investigate the effects of the observers’ responses on their EEG activities, to uncover the EEG signatures reflecting the perception of trustworthiness in real-time social contexts.

## 2. Materials and Method

### 2.1. Description of the Dataset

As illustrated by Chen, Fazil, and Wallraven (2024), the dataset we used in the present investigation was curated from a data description study published in *Scientific Data.* This dataset contains a complete set of raw behavioral data and EEG data, as well as a preprocessed EEG dataset that were collected in a two-player deception experiment. Twenty-four participants (12 females, age: 25 ± 4.34 yrs) with normal or corrected-to-normal vision and no reported history of neurological disorder participated in this experiment for monetary rewards of around USD10/h. This experiment was approved by the Institutional Review Board with the number KUIRB-2019-0043-01. All participants were naïve to the experimental paradigm and gave written informed consent prior to the experiment.

### 2.2. Experimental Procedures

Depicted in Figure 1, the task of this experiment involved two participants serving as the player and the observer, respectively, at each time. The two participants faced each other and sat in front of two separated 24-inch monitors (resolution: 1920×1080 px^2^, fresh rate: 60Hz; produced by LG, South Korea), with a HD pro C920 webcam (Logitech, Switzerland) installed on the top of the player’s monitor (Figure 1 b). Schematic in Figure 1 a, a trial began with a fixation period for 1000ms, during which both participants were asked to fixate on a fixation cross appeared in the center of their respective monitors. Then, a card contained a colored digit (ranging from 1 to 6) presented on the monitor of the player’s side, whereas on the observer’s side, the monitor showed the live stream of the player’s face. This stage lasted for 3000ms, and during this period, the player had to decide what number to be relayed to the observer depending on the color (black, purple, or blue) of the presented digit. The three colors cued three different strategic conditions: (1) the instructed truth condition that the player had to relay the same number presented in the card to the observer; (2) the instructed lie condition that the player had to relay a different number than the presented one; and (3) the spontaneous condition that the player chose any number within the range 1-6 to relay. Succeeding this player decision stage, the player was asked to relay the number by pressing a correspondent button on a RB-740 response pad (Cedrus Corporation, USA) within 3000ms, during which only the background was presented on the observer’s side. Triggered by the player’s response, a card showing the relayed number in black presented on the observer’s monitor, and the observer was asked to decide whether the information was a lie (perceived untrustworthy) or the truth (perceived trustworthy) by pressing one of two buttons on the response pad within 3000ms. Upon the observer’s response, feedback was given to both participants according to a score system as illustrated in Figure 1 c. The score system, explained before the experiment, was designed to encourage active participation of both participants by successful deceiving and detecting in exchange for higher scores. At the end of each trial, for the monitors of both sides, the trial scores that both participants acquired were presented for 1000ms and followed by a summary of status showing the accumulated scores, number of trials and rounds won by each side, as well as the progress of experiment for 3000ms.

**Figure 1.**
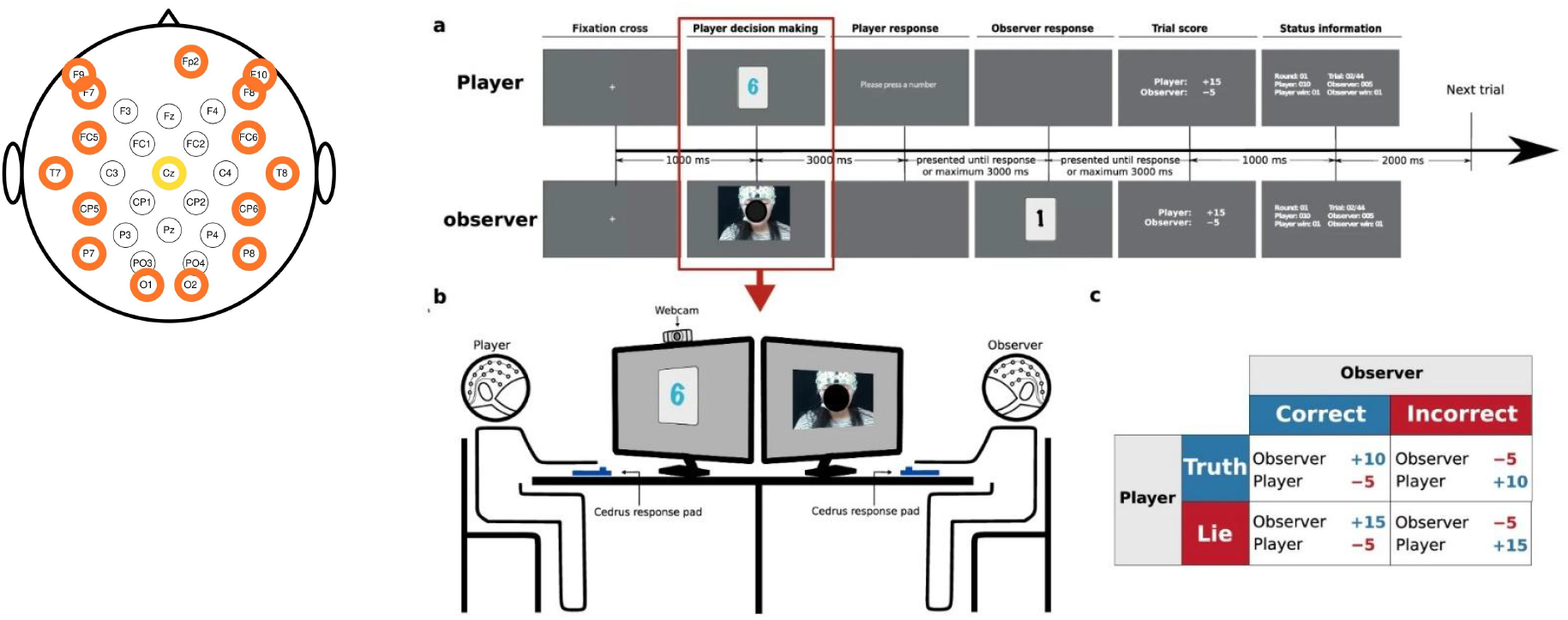
Illustration of the experiment. (a) Schematic of a trial. (b) Setup of the experiment. (c) Scoring system used in the experiment. This figure was copied from Figure 1 in Chen, Fazil, and Wallraven (2024).

The 24 participants were separated into 12 pairs, and in each pair, two participants took turns to serve as the player and the observer. This resulted to 24 paired cases, and 23 of them successfully finished the experiment. For each case, the entire experiment consisted of 11 sessions, in which each session contained 44 trials (spontaneous: 22, instructed lie: 11, instructed truth: 11) and these trials were presented in a randomly shuffled order. During all these experiments, EEG and responses of both participants were recorded. This experiment was programmed in Python using PsychoPy.

### 2.3. EEG Apparatus

According to the data description, the EEG data was recorded with two BrainAmp amplifiers (Brain Products, Germany) from 30 EEG electrodes and an EOG electrode. Electrode locations on the caps are visualized in Figure 1. Due to connection issues, the Oz channel was removed from the dataset. The data was digitized at 500Hz, nose-referenced online, and forehead grounded to the electrode Fpz.

### 2.4. Data Analysis

#### 2.4.1. Data Preprocessing

In the preprocessed dataset, the data was down-sampled to 100Hz and filtered at 1-49Hz. Artifacts were detected and rejected using EEGLAB functions and ICA-based algorithms with Matlab (The MathWork, USA). Data from each channel was then baseline corrected by 500ms prior to the stimulus onset. Further details about the algorithms during preprocessing can be found in the data description (Chen, Fazil, and Wallraven, 2024). Since this study targeted on facial trustworthiness, we selected the observers’ data from the “player decision making” stage (as labeled in Figure 1 a), which lasted for 3500ms beginning from 500ms prior to the facial expression display, and ending 3000ms afterwards. The resulted EEG signals were then re-referenced to the electrode Cz.

We conducted four analyses subjective to the observers’ responses, either trustworthy or untrustworthy, toward the players. In these analyses, we analyzed the effects of the observer’s perception of trustworthiness by examining how their EEG data could reflect observers’ responses. First, we performed an ERP analysis to examine whether these social evaluations elicited amplitude differences across scalp electrodes and to identify the associated cortical regions. Second, to explore hemispheric asymmetries, we paired electrodes across the left and right hemispheres and compared the magnitude of their signals, assessing how these interhemispheric differences were modulated by the trustworthiness perception. Third, we extracted the instantaneous phases of EEG signals and repeated the interhemispheric analysis based on phase differences between homologous electrode pairs, enabling a finer-grained investigation of cortical synchrony and temporal alignment related to trustworthiness perception. A linear regression was performed on the phase progression related profiles with respect to time to reflect the general frequency of the original signals. Finally, we calculated the weighted average of the frequencies of selected signals in a spectral centroid analysis to further examine how the instantaneous phase was affected.

#### 2.4.2. ERP Analysis

For each observer, we segmented the EEG data from into two conditions based on their responses: one comprising trials in which the observer perceived trustworthy from the player, and the other comprising trials in which the observer perceived untrustworthy. We first examined the behavioral and neural strategies by analyzing ERPs. Subsequently, we identified electrodes and time windows that exhibited signal differences associated with the observer’s responses. This was done by averaging ERP signals across all trials and participants separately. To assess the statistical significance of these effects, we conducted repeated-measures ANOVAs (RM-ANOVAs) on the mean signal amplitudes within individuals, with observer attitude (trustworthy versus untrustworthy) as the within-subject factor. In order to test the effects from trustworthiness against other factors, we grouped the data by observers’ correctness (correct verses incorrect), and by random segmentation (denoted by “control”) in each epoch that were balanced among the conditions to be tested.

#### 2.4.3. Interhemispheric Electrode Pairs

To examine interhemispheric differences, we paired electrodes across the frontal-posterior hemispheres, selecting those located at symmetric scalp sites. This resulted in twelve electrode pairs: F3–F4, F7–F8, F9–F10, FC1–FC2, FC5–FC6, C3– C4, CP1–CP2, CP5–CP6, P3–P4, P7–P8, PO3–PO4, and O1–O2. For each pair, we computed difference waveforms by subtracting the signal from the right hemisphere electrode from its left hemisphere counterpart. Consistent with the ERP analysis, EEG signals from each participant were divided into two conditions based on the observer’s responses. We then averaged the signals within each condition to generate signal profiles for statistical analysis.

#### 2.4.4. Instantaneous Phase Analysis

The instantaneous phase dynamics of a signal was computed using the approach of Hilbert transformation, as illustrated by equations 1 and 2.

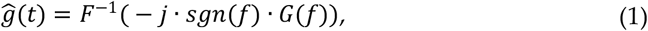

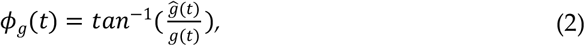

where ĝ (t) is the Hilbert transform of a given time-series g(t), F^−1^ is the inverse Fourier transform of the input signal, G(f) is the Fourier transform of g(t), and ϕ_g_(t) is the instantaneous phase of g(t). We then unwrapped *ϕ_g_* (*t*) to remove the discontinuity of the instantaneous phase profile and uncover the phase progression.

#### 2.4.5. Spectral Centroid Analysis

The spectral centroid of a EEG signal was computed using equation 3 based on the its power spectral density (PSD) ranging from 0 to 50 Hz along the frequency domain with a bin of 0.5 Hz.

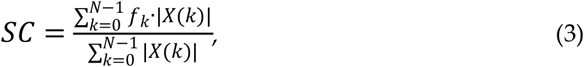

where SC stands for spectral centroid, f_k_ and |X(k)| are frequency and magnitude of PSD at bin k.

## 3. Results

### 3.1. Behavioral Results

For the 484 trials each participant ran as an observer, the mean (std) number of correct and incorrect trials were 241.13 (13.56) and 242.74 (13.5), respectively. The number of trials, in which observers’ perceived trustworthiness by two choices, were 244.91 (10.63) for trustworthy and 238.96 (10.64) for untrustworthy. As for the random control condition, each participant’s trials were segmented into two groups with counts of 242.13 (0.74) and 241.74 (0.85). Figure 2 illustrates how the count of trials varied by these conditions. A three-factor RM-ANOVA with the correctness, trustworthiness, and control as the main factors on the count of trials found no significant effect of correctness (F(1,22)=0.08, p=0.78), no significant effect of trustworthiness (F(1,22)=1.8, p=0.19), and no significant effect of control (F(1,22)=1.45, p=0.24). Moreover, the control condition was not significantly interacted with correctness (F(1,22)=0.43, p=0.52) and trustworthiness (F(1,22)=0.63, p=0.44), while the interaction between correctness and trustworthiness was significant (F(1,22)=7.29, p=0.01).

**Figure 2.**
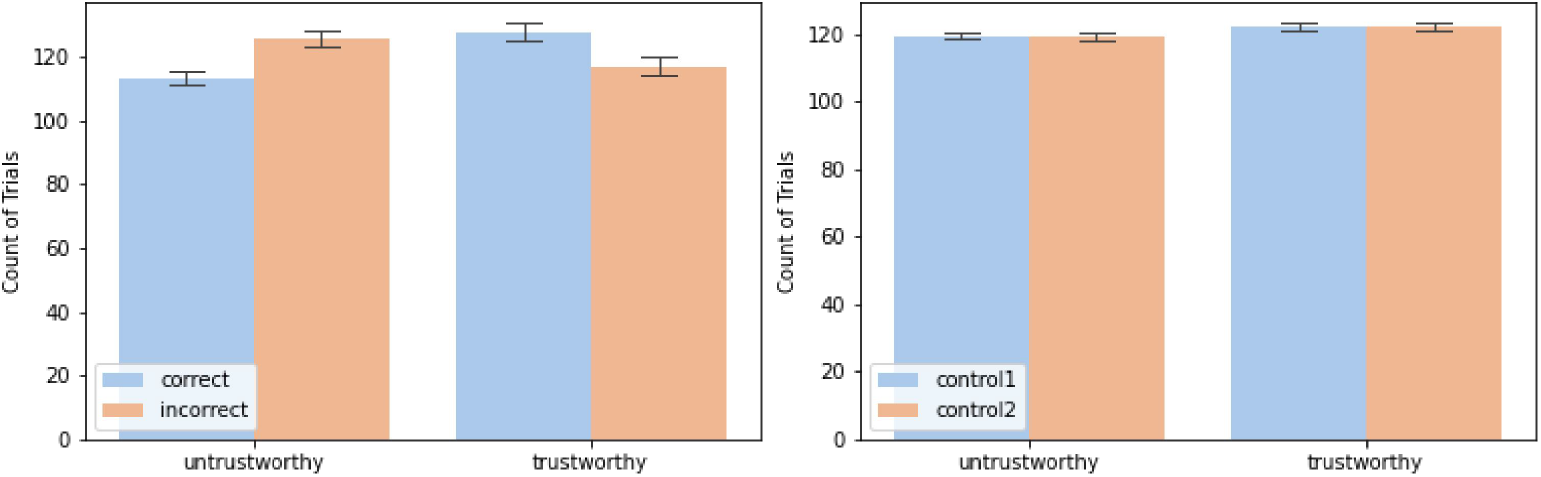
Count of trials by trustworthiness (trustworthy versus untrustworthy), correctness (correct versus incorrect), and random segment control (control1 versus control2).

We found the observers’ performance was not significantly differed than the chance level and their perceived trustworthiness by two choices were also not significantly uneven. Although the interaction between trustworthiness and correctness suggests that observers’ judgments was not random, they were largely inaccurate, indicating the reliance on intuitive processing. These findings are consistent with current theories suggesting that face-based social evaluations are intuitive and inherently challenging without additional contextual information (see review by Todorov et al., 2015). Regarding the random control condition, we observed a balanced distribution of counts across both trustworthiness and correctness, supporting its validity as a baseline for comparing against potential random effects.

### 3.2. ERP Analysis

Figure 3 presents the ERPs recorded from frontal and posterior electrodes (Fz, P7, P8, O1, and O2). A clear N170 component was detected, as shown in the posterior signals in Figure 3 that a dip at around 200ms between two adjacent positive peaks, which is a typical ERP signature, usually peaking between 140 and 230ms, linked to the perception of emotional face-body compound stimuli (Meeren et al., 2005; Blau et al., 2007; Hinojosa et al., 2015). This finding underscore the observers’ engagement that focusing on interpreting the facial expressions of the players.

**Figure 3.**
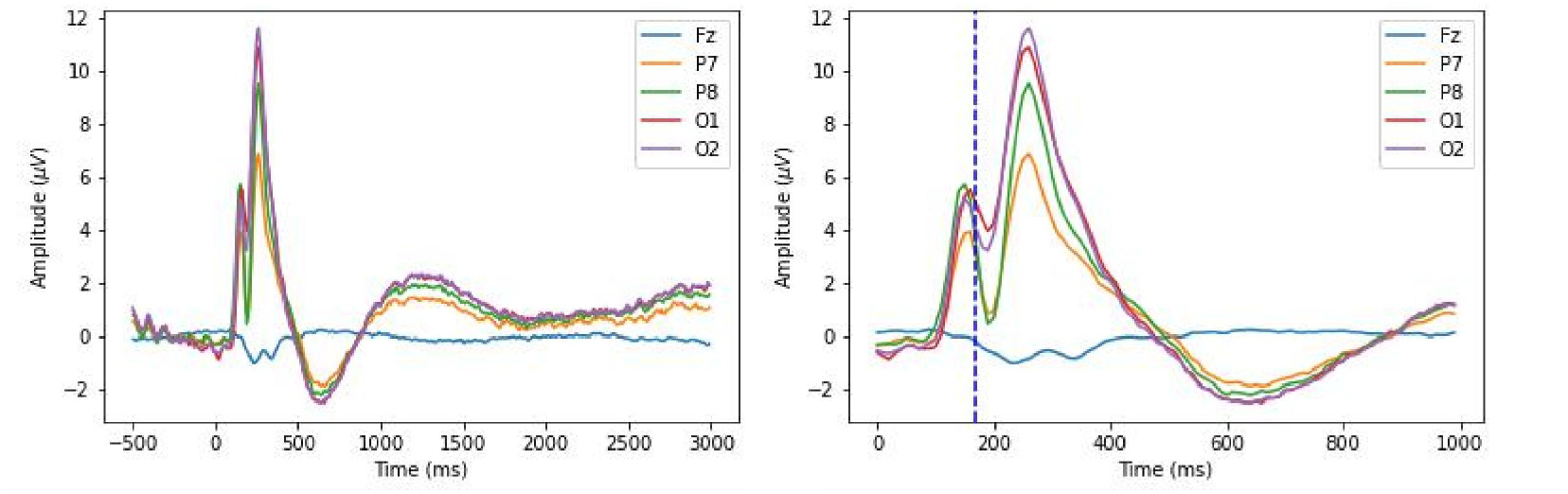
Grand average ERPs for the observers. The left panel shows the signals during the entire trial. The right panel shows the signals during 1000ms post-stimulus, and a dashed line marks 170ms.

However, we didn’t find any trustworthiness effects on the magnitude of EEG signals. Figure 4 presents the topographical distribution across the cortex of the difference in EEG signals between trustworthy and untrustworthy trials. Each graph visualizes averaged ERP signals across 100ms. We found the grand difference between trustworthy and untrustworthy related trials across all the electrodes were below 1*μV*, and RM-ANOVAs detected no significant effects from any duration.

**Figure 4.**
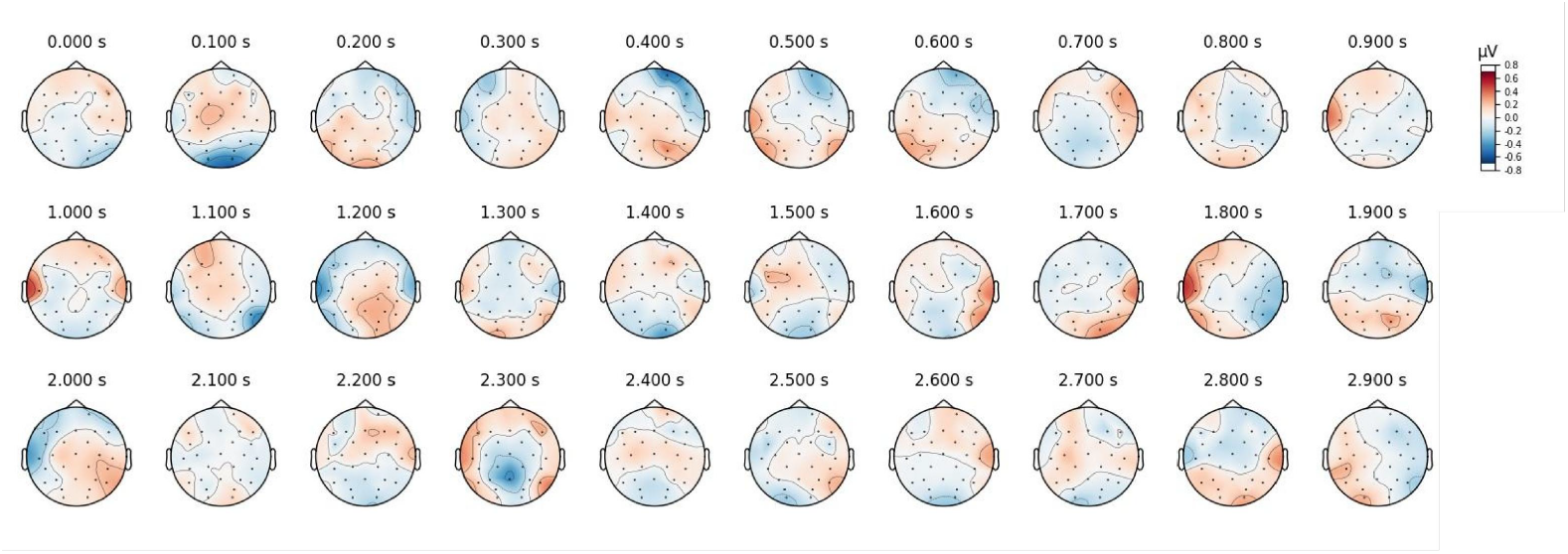
Topographics of grand ERP difference between trustworthy and untrustworthy epochs.

### 3.3. Interhemispheric Comparisons

#### 3.3.1. Interhemispheric Difference in Signal Magnitude

Figures 5 illustrates the pairwise signal differences between electrodes in the left and right hemispheres, categorized by trustworthy, untrustworthy, correct, incorrect, and two control conditions. We observed that two peaks before and after the N170 component were stronger in the right hemisphere compared to the left, mostly evident in P7-P8, where subtracting P8 from P7 leads to negative peaks. This resulted in a larger N170 component in the right hemisphere, which can also be detected in Figure 3. This right-lateralized N170 effect in face-selective tasks has been consistently reported in previous studies (Ibáñez et al., 2012; Nakajima et al., 2012; Gao et al., 2019), especially in differentiating between word- and face-related N170 components (Maurer et al., 2008; Rossion et al., 2003; Dundas et al., 2014). These findings confirmed the involvement of emotional face recognition of observers during experiment.

**Figure 5.**
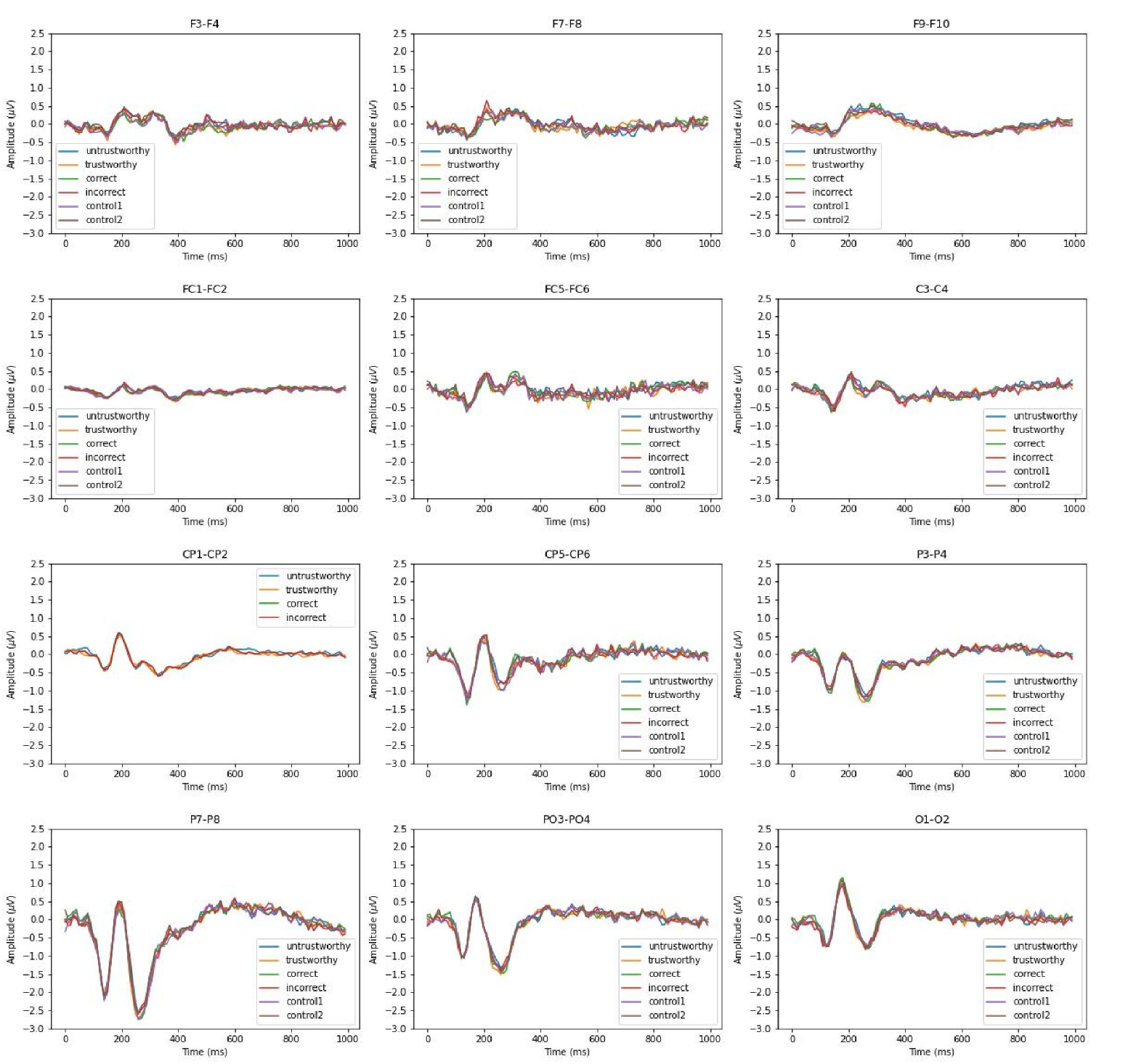
Grand average interhemispheric ERP differences conditioned by trustworthy, untrustworthy, correct, and incorrect.

As for the interhemispheric effects comparing all the conditions, we found these profiles of pairwise difference indifferent than each other for every electrode pair. RM-ANOVAs confirmed the absence of significant effects for any electrode pair during any of the major time windows.

#### 3.3.2. Interhemispheric Difference in Signal Phase Progression

In this section, we synthesized the phase progression of the signals from each electrode and calculated the phase shift between interhemispheric electrode pairs. To examine the effect of trustworthiness, we determined the *conditioning interhemispheric phase shift*, by calculating the absolute difference in interhemispheric phase shift between trustworthy- and untrustworthy-related epochs, and we controlled it by that between correct- and incorrect-related epochs, as well as two random control groups. Our null hypothesis was that these three profiles of *conditioning interhemispheric phase shift* representing trustworthiness, correctness, and random control should be equal if trustworthiness can induce no effect on the interhemispheric phase shift progression.

Figure 5 illustrates the effects of trustworthiness on interhemispheric phase shifts across electrode pairs. We observed that phase dynamics were generally asymmetrical between the two hemispheres. Also, these effects appeared to be highly variable, as *conditioning interhemispheric phase shift* profiles on all three conditions deviated from zero over time across electrode pairs.

Comparing phase profiles among the three conditions, significant effects were observed in the central and parietal regions, most notably at the CP1–CP2 electrode pair from 1500 ms to 3000 ms. A RM-ANOVA detected a significant effect on the *conditioning interhemispheric phase shift* during this interval (F(2, 44) = 9.37, p<0.001, 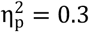). Pairwise comparison tests (Tukey’s HSD) detected a significant difference between trustworthiness and control (p=0.003), as well as between trustworthiness and correctness (p=0.003), but failed to find any significant difference between correctness and control (p=0.9). Considering that instantaneous phase progression reflects how the phase of an oscillatory EEG signal evolves over time, these results suggest that such phase dynamics become more asymmetrical in two hemispheres during facial trustworthiness perception than random states.

From equations 1 and 2, we know that the instantaneous phase is the integral of instantaneous frequency over time. We thereby performed linear regressions on each participant’s data at CP1 - CP2 and analyzed the general frequency of their EEG signals based on the resulted slopes. Figure 7 shows the results of linear regressions and the slopes in a unit of Hz are illustrated in Figure 8. The average frequency difference (slope) conditioned by trustworthiness, correctness, and control were 2.18 ± 2.16 Hz, 1.02 ± 1.1 Hz, and 0.99 ± 0.74 Hz, respectively. A RM-ANOVA comparing the these conditions confirmed a significant effect on the slopes (F(2, 44) = 5.14, p<0.01, 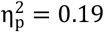). Pairwise comparison tests (Tukey’s HSD) detected a significant difference between trustworthiness and control (p=0.02), as well as between trustworthiness and correctness (p=0.023), but failed to find any significant difference between correctness and control (p=0.9). These results indicate a higher divergence between two hemispheres by means of signal frequency induced by facial trustworthiness perception than random states.

More specifically, the frequency difference between signals in CP1 and CP2 was 0.99 Hz by average under different random states, but perceiving trustworthy and untrustworthy faces induced different interhemispheric divergence with a degree (2.18 Hz by average) significantly higher than random levels.

### 3.4. Spectral Centroid Analysis on CP1 - CP2

In the instantaneous phase analysis, we detected major effects induced by facial trustworthiness perception on the phase difference between CP1 and CP2, which was tested to be significant against the random control. Subsequent linear regression analysis determined diverging speeds in phase progression underlying such phenomena, which indicates different levels of hemispheric asymmetry in signal frequency associated with trustworthy versus untrustworthy perception. To further investigate how these signals from two hemispheres different than each other in terms of frequency and thereby reflect the effects of facial trustworthiness perception, we computed the spectral centroid of the signals at the electrode pair CP1 - CP2. Eye observations from the profiles of phase shifts in Figure 6 suggested high slope difference between trustworthiness and control conditions during 800 - 2000 ms. We therefore selected the EEG signals within this period and performed the spectral centroid analysis on them conditioned by trustworthy, untrustworthy, correct, incorrect, and two random controls.

**Figure 6.**
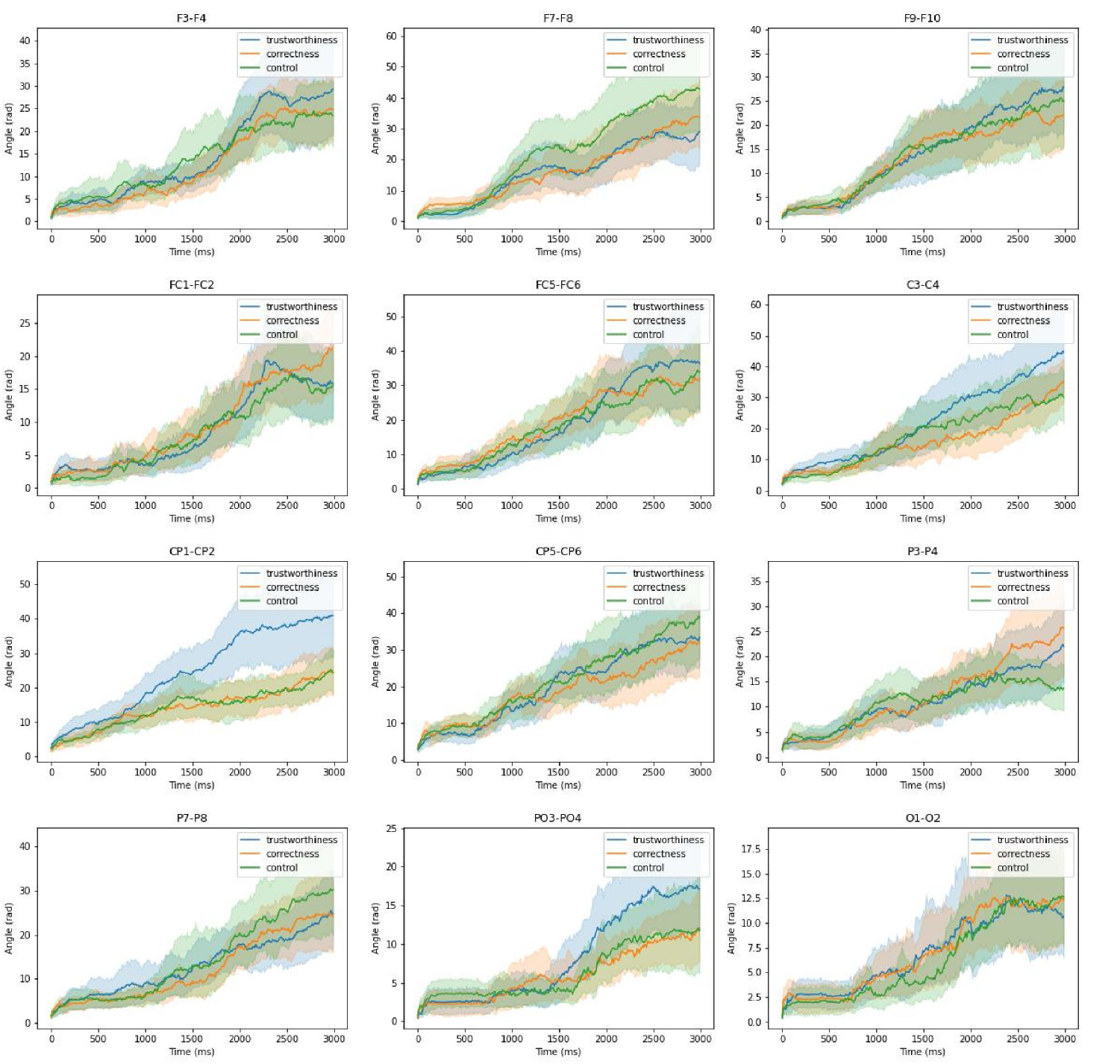
Profiles of conditioning interhemispheric phase shift on trustworthiness, correctness, and control. The shadings indicate 95% confidence intervals across participants.

**Figure 7.**
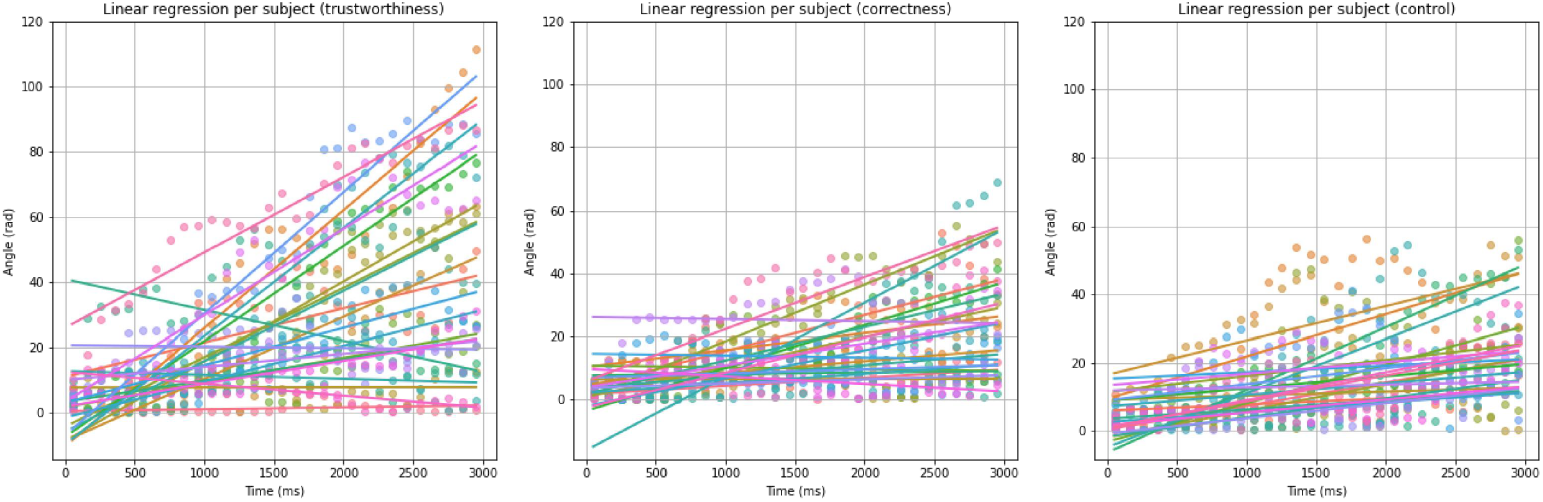
Results of linear regression per subject for trustworthiness, correctness and control conditions.

**Figure 8.**
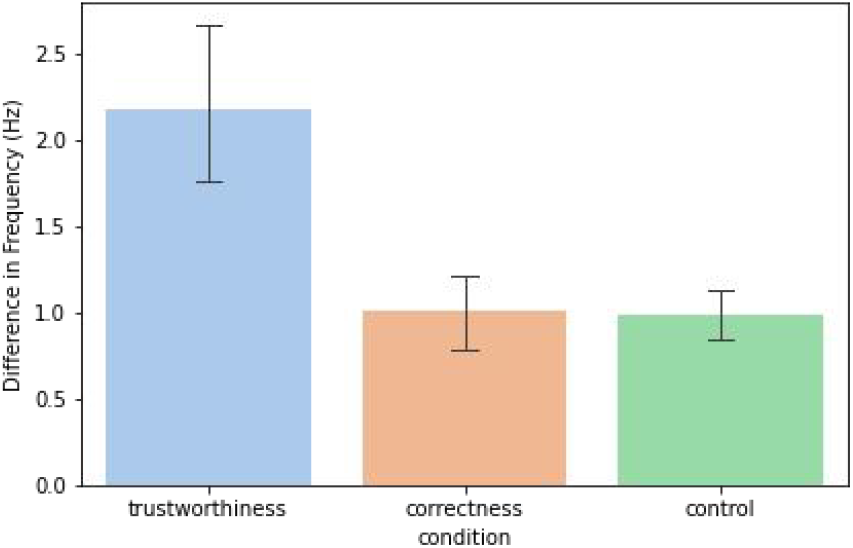
Slopes resulted by the linear regression analysis as transformed to be in a unit of Hz.

As shown in Figure 9, we found that the spectral centroid at CP1 - CP2 was dependent on trustworthiness. A RM-ANOVA with two factors (channel and trustworthiness) detected a significant interaction between these two factors (F(1, 22) = 6.44, p = 0.018, 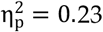), but didn’t detect any factorial effect (channel: F(1,22) = 0.14, p = 0.71,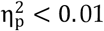; trustworthiness: F(1,22) = 0.03, p = 0.86, 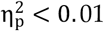). Such interaction was absent when the data was conditioned by correctness (channel: F(1,22) = 0.46, p = 0.5, 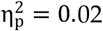; correctness: F(1,22) = 0.004, p = 0.94,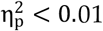; channel×correctness: F(1,22) = 0.004, p = 0.94,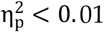) or by control (channel: F(1,22) = 0.001, p = 0.975,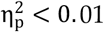; control: F(1,22) = 0.014, p = 0.91,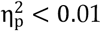; channel×correctness: F(1,22) = 0.052, p = 0.82, 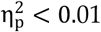).

**Figure 9.**
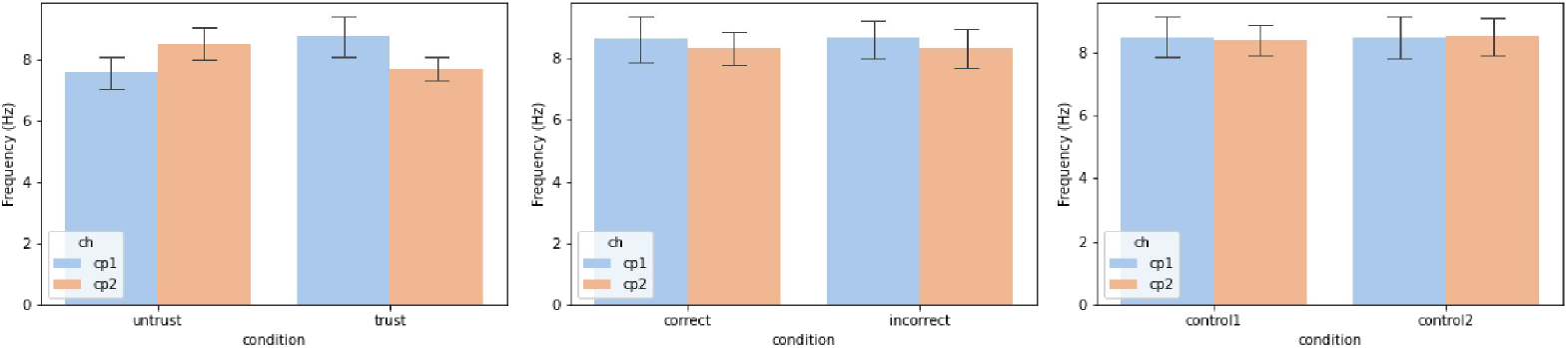
Spectral centroid of the signals from the electrode pair CP1 - CP2.

These results further support our findings from the instantaneous phase analysis, indicating that hemispheric asymmetry in the centroparietal regions reflects the perception of facial trustworthiness. Specifically, the observed phase asymmetry appears to be driven by a lateralization of signal frequency: trustworthy faces were likely to induce relatively higher frequencies in the left hemisphere, while untrustworthy faces could induce relatively higher frequencies in the right hemisphere.

## 4. Discussion

In this study, we analyzed EEG data to investigate how observers evaluate facial trustworthiness during deceptive behavior. Unlike previous neuroimaging research that primarily used static facial images, this study examined neural activity in a naturalistic setting, where observers viewed live video streams of players’ faces in real time. During the deception-encouraging game, observers were actively engaged, as reflected in both their behavioral responses and the presence of the N170 component in their ERPs. Furthermore, we found that the perception of facial trustworthiness was dynamically modulated by lateralized brain activity in the observers’ centroparietal cortex, with distinct patterns emerging between the left and right hemispheres between 1500-3000ms post-stimulus. Whereas earlier studies using static face stimuli suggested that facial trustworthiness perception is rapid and prolonged exposure to facial stimuli did not substantially alter trustworthiness judgments, our findings emphasize the role of dynamic and persistent affective processing in real-time social interactions.

In this study, we found that participants’ behavioral responses were balanced across both trustworthiness and correctness, aligning with previous research suggesting that judgments of facial trustworthiness can be intuitive and often inaccurate (Bond et al., 2006; Bond & DePaulo, 2008; Rule et al., 2013; see review by Todorov et al., 2015). One possible explanation is the inherent instability of facial appearance. Prior studies have shown that participants exhibit high variability when identifying faces across different photographs of the same individual, comparable to the variability observed between different individuals (Jenkins et al., 2011; Burton et al., 2016). This high degree of variation has also been observed in social attribution tasks, where judgments about traits like trustworthiness showed similar within- and between-individual variability (Olivola & Todorov, 2010). Although often inaccurate, research indicates that trustworthiness judgments can be made extremely quickly, within the first 100 ms of stimulus presentation (Willis & Todorov, 2006), a finding supported by EEG studies showing early ERP components reflecting such evaluations (Yang et al., 2011).

However, our findings indicate that when viewing faces in real-time, trustworthiness perception is dynamic, as reflected by a sustained interhemispheric phase divergence in the centroparietal cortex, driven by a lateralization effect in EEG frequency. Emotion-related hemispheric lateralization has been widely documented in affective neuroscience (see review by Harmon-Jones et al., 2009). Early studies reported greater alpha activity lateralized to the left hemisphere in affective conditions, particularly in posterior regions (Smith et al., 1987). Furthermore, associations have been found between left frontal electrical activity and emotional facial expressions, with increased activation observed during joyful expressions (Ekman & Davidson, 1993; Graham & Cabeza, 2001). Asymmetrical neural activity has also been linked to motivational direction: for instance, promotion-focused states have been associated with stronger bilateral frontal asymmetries than prevention-focused states (Amodio et al., 2004). Taken together, our findings suggest that the evaluation of facial trustworthiness in naturalistic, real-time settings is influenced by a persistent and dynamic affective modulation, possibly rooted in these well-established lateralization mechanisms..

Another insight of this investigation is to analyze instantaneous phase of EEG signals to uncover the neural dynamics underlying human behaviors in naturalistic settings. Analyzing ERPs in real-time, naturalistic scenarios poses significant challenges due to the complexity and variability of uncontrolled environments. Traditional ERP analyses often rely on averaging across many trials under controlled conditions to isolate stimulus-locked brain responses. However, in dynamic, ecologically valid settings, such as face-to-face interactions in this investigation, the timing of neural responses can vary significantly, making it difficult to align and interpret ERP components. In this study, viewing actual faces in a real-time manner induced different ERP patterns compared to those of previous studies using static facial stimuli. Also, magnitude-based measures failed to detect any trustworthiness effects, but examining the instantaneous phase of neural oscillations offers a promising alternative. Previous studies have demonstrated the effectiveness of this approach in uncovering neural dynamics involved in auditory-related cognitive functions, such as speech comprehension (Li et al., 2022) and music perception (Poikonen et al., 2018). The present study extends this evidence by showing that it can also reveal the neural underpinnings of vision-based cognitive processes.

## Author Contributions

Conceptualization, H.Q. and D.H.; methodology, H.Q. and D.H.; software, D.H.; validation, H.Q. and D.H.; formal analysis, H.Q. and D.H.; investigation, H.Q. and D.H.; resources, D.H.; data curation, D.H.; writing—original draft preparation, H.Q. and D.H.; writing—review and editing, D.H.; visualization, H.Q. and D.H.; supervision, D.H.; project administration, D.H.; funding acquisition, D.H. All authors have read and agreed to the published version of the manuscript.

## Data Availability Statement

The data used in the paper is readily accessible on Figshare at https://figshare.com/articles/dataset/An_EEG_Dataset_of_Neural_Signatures_in_a_Competitive_Two-Player_Game_Encouraging_Deceptive_Behavior/24760827, with a CC BY 4.0 license.

